# Therapeutic Targeting of 15-PGDH in Murine Idiopathic Pulmonary Fibrosis

**DOI:** 10.1101/2019.12.16.878215

**Authors:** Julianne N.P. Smith, Matthew D. Witkin, Alvin P. Jogasuria, Kelsey F. Christo, Thomas M. Raffay, Sanford D. Markowitz, Amar B. Desai

**Author notes:** Denotes co-corresponding author. Correspondence: Amar B. Desai, Department of Medicine, Case Western Reserve University.

## Abstract

Idiopathic pulmonary fibrosis (IPF) is a progressive disease characterized by interstitial remodeling and pulmonary dysfunction. The etiology of IPF is not completely understood but involves pathologic inflammation and subsequent failure to resolve fibrosis in response to epithelial injury. Therapeutic strategies for IPF are limited to anti-inflammatory and immunomodulatory agents, which are only partially effective. Prostaglandin E2 (PGE2) disrupts TGFβ signaling and suppresses myofibroblast differentiation, however practical strategies to raise tissue PGE2 during IPF have been limited. We previously described the discovery of a small molecule, (+)SW033291, that binds with high affinity to the PGE2-degrading enzyme 15-hydroxyprostaglandin dehydrogenase (15-PGDH) and increases PGE2 levels. Here we evaluated pulmonary 15-PGDH expression and activity and tested whether pharmacologic 15-PGDH inhibition (PGDHi) is protective in a mouse model of bleomycin-induced IPF. Long-term PGDHi was well-tolerated, reduced the severity of pulmonary fibrotic lesions and extracellular matrix remodeling, and improved pulmonary function in bleomycin-treated mice. Moreover, PGDHi attenuated both acute inflammation and weight loss, and decreased mortality. Endothelial cells and macrophages are likely targets as these cell types highly expressed 15-PGDH. In conclusion, PGDHi ameliorates inflammatory pathology and fibrosis in murine IPF, and may have clinical utility to treat human disease.

**Article summary:** In IPF, lung epithelial injury leads to local inflammation and fibrosis, which impairs pulmonary function. Inhibition of 15-PGDH using a well-tolerated small molecule attenuates inflammation and prevents pulmonary fibrosis and dysfunction in a mouse model of bleomycin-induced IPF.

## Introduction

Idiopathic pulmonary fibrosis (IPF) is a progressive and irreversible disease involving the accumulation of extracellular matrix throughout alveoli and interstitial spaces, leading to the destruction of lung parenchyma and impaired gas exchange ^1^. Although the precise etiology of IPF is unknown, the median age of onset is 66 ^2^, likely due to dysfunctional wound healing ^3^, heightened inflammation, and a reduced ability to resolve fibrosis ^4^ as an organism ages. Anti-fibrotic agents are commonly used to treat IPF but show limited efficacy, as evidenced by the 3-4 year estimated survival in IPF patients ^2^. Therefore, novel therapeutic approaches to prevent IPF pathogenesis and mitigate IPF severity are needed.

The initiation of IPF is often characterized by early recurrent lung epithelial injuries that are not cleared and eventually lead to the deposition of fibrosis. Although anti-inflammatory therapies have provided little benefit in IPF trials ^5^, studies suggest that immune responses are involved in disease development and progression ^6^. Indeed, neutrophils accumulate in IPF patient lungs ^7-9^, and neutralizing neutrophil-derived products mitigates murine IPF severity ^10,11^. Monocyte-derived alveolar macrophages are also implicated in human IPF ^12^. Notably, classically-activated, or M1, macrophages produce potent pro-inflammatory cytokines including TNFα, IL-1, and IL-6 ^13^. In response to chronic inflammation, M1 macrophages take on characteristics of alternatively-activated, or M2, macrophages ^14^, which contribute to fibrosis and collagen synthesis via production of transforming growth factor-β (TGFβ), platelet-derived growth factor (PDGF), and upregulation of L-arginine metabolism ^13^.

Fibroblasts play a key role in IPF pathology through their proliferation, migration, and differentiation to myofibroblasts. TGFβ drives the differentiation of fibroblasts to myofibroblasts via the induction of various cellular processes and signaling cascades, including the activation of Smad proteins, phosphatidylinositol 3-kinase/Akt, ERK and MAPK signaling, and cytosolic calcium oscillation ^15,16^. Engagement of these signaling pathways results in the expression of ECM proteins, and the formation of stress fibers. Several lung-resident and circulating cell types produce PGE2 upon inflammation, and specifically following bleomycin-induced lung injury, including pulmonary fibroblasts, alveolar epithelial cells, and monocyte/macrophage lineage cells ^17,18^. Signaling via PGE2 receptors EP2 and EP4 antagonizes TGFβ-induced pro-fibrotic signaling, suggesting that this axis is critical for normal pulmonary tissue repair and regeneration. Endogenous PGE2 production is likely insufficient to block pathogenesis due to the reduced expression of EP2 and EP4 in fibrotic lung tissue ^19,20^, however. Therefore, well-tolerated strategies to increase pulmonary PGE2 levels are likely to mitigate pathogenesis. Indeed, recent reports have demonstrated that either systemic administration of the long-acting PGE2 analog 16,16-dimethyl-PGE2 ^18^, or targeted PGE2 delivery via pulmonary endothelial cell-specific antibodies or inhalation of PGE2-loaded liposomes ^21,22^ demonstrate therapeutic efficacy in murine pulmonary fibrosis. *In vitro*, PGE2 stimulation or EP2/EP4-specific agonism, abrogates myofibroblast differentiation and expression of ECM genes in TGFβ-treated human pulmonary fibroblasts and in fibroblasts derived from IPF patients ^19,23-25^. Thus, we hypothesized that increasing endogenous PGE2 by systemic administration of well-tolerated small molecules that inhibit the PGE2-degrading enzyme 15-hydroxyprostaglandin dehydrogenase (15-PGDH) would prevent pulmonary fibrosis in bleomycin-treated mice.

## Results

### 15-PGDH is highly expressed and active in healthy murine lung tissue

(+)SW033291 is known to increase systemic PGE2 levels and enhance tissue regeneration ^26^. To determine if 15-PGDH may be targetable in the murine lung, we first examined its expression in the lungs of healthy mice, relative to other organs in which 15-PGDH inhibitors (PGDHi) have demonstrated therapeutic efficacy ^26,27^. Immunohistochemical staining revealed subsets of PGDH+ hematopoietic cells in the BM and numerous PGDH+ cells in the colonic epithelium (**Fig. 1A-B**). In contrast, 15-PGDH was highly expressed throughout the lung parenchyma (**Fig. 1C**), suggesting pulmonary tissue may also be responsive to PGDH inhibition. We further compared expression of *Hpgd*, the gene that encodes 15-PGDH, in colonic and pulmonary tissue relative to BM. Lung tissue homogenates displayed 25- and 3-fold higher *Hpgd* gene expression than BM and colon, respectively (**Fig. 1D**). To confirm that pulmonary 15-PGDH is functional and therefore capable of regulating local PGE2 levels, we next measured specific enzymatic activity in homogenates from the same organs. Lung tissue demonstrated >200 and 3.7 fold higher activity per milligram total protein than BM and colonic tissue, respectively (**Fig. 1E**). Together these data demonstrate that 15-PGDH is abundantly expressed and highly enzymatically active in the murine lung and provide rationale for pharmacologic targeting of 15-PGDH as a strategy to increase local PGE2 and reduce fibrosis in IPF.

**Figure 1:**
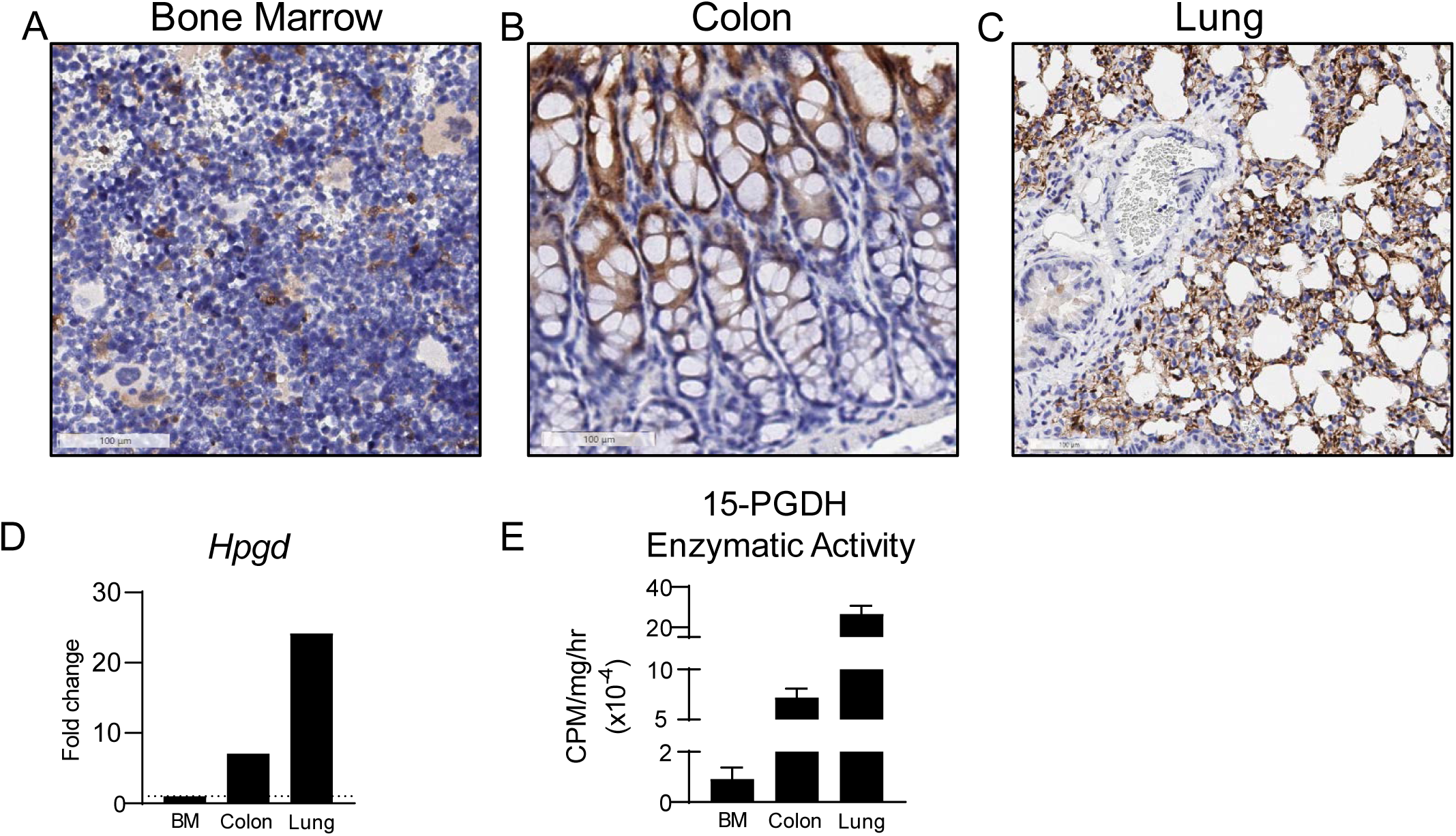
15-PGDH is highly expressed in the murine lung. (**A-C**) Representative images of 15-PGDH staining (brown) in healthy murine bone marrow (BM), colon, and lung, with Hematoxylin counter stain. 20X, scale bars represent 100μm, as indicated. (**D**) Relative gene expression of *Hpgd* in murine BM, colon, and lung by RT-PCR, normalized to *B2m* levels and expressed as fold change relative to the level in BM. (**E**) 15-PGDH enzymatic activity in murine BM, colon, and lung, as measured in counts per minute (CPM) over one hour and normalized to input protein (in mg).

### PGDHi mitigates early bleomycin-induced inflammation

In mice, administration of bleomycin results in lung injury that mimics key aspects of human IPF ^28^, with an initial inflammatory phase followed by a subsequent fibrotic phase ^29^. To study the effects of 15-PGDH inhibition in IPF, we administered intravenous bleomycin and began twice daily treatment of mice with (+)SW033291 (PGDHi) or vehicle control (**Fig. 2A**). PGDHi attenuated early pulmonary inflammation, as evidenced by greater than 50% reductions in *Il1b* and *Il6* expression in lung tissue 7 days post-bleomycin exposure, in addition to moderate reductions in the expression of other inflammation-associated genes (**Fig. 2B**). These data indicate that inhibiting 15-PGDH in the context of bleomycin-induced lung injury may limit pathologic inflammation in the lung. Moreover, PGDHi treatment was associated with significantly lower levels of the neutrophil chemoattractant KC(CXCL1)/GRO, the cytokine TNFα, and a trend towards reduced IL-10 in the serum (**Fig. 2C**). TNFα is known to promote TGFβ1 expression ^30^, and although IL-10 limits inflammation in many contexts, it is thought to promote fibroblast proliferation in IPF ^31^, therefore these changes likely have an anti-fibrotic effect on injured pulmonary tissue. Of note, TGFβ1 was elevated in the serum 7 days post-bleomycin exposure and PGDHi treatment led to a moderate but statistically insignificant reduction compared to vehicle (44±5.5 versus 36±1.9 pg/mL). Additionally, while vehicle-treated mice displayed alveolar wall thickening and an abundance of small alveolar spaces, a trend towards greater alveolar size was observed with PGDHi (**Fig. 2D**). These data therefore indicate that PGDHi limits both systemic and local inflammation during the early phase of bleomycin-induced lung injury.

**Figure 2:**
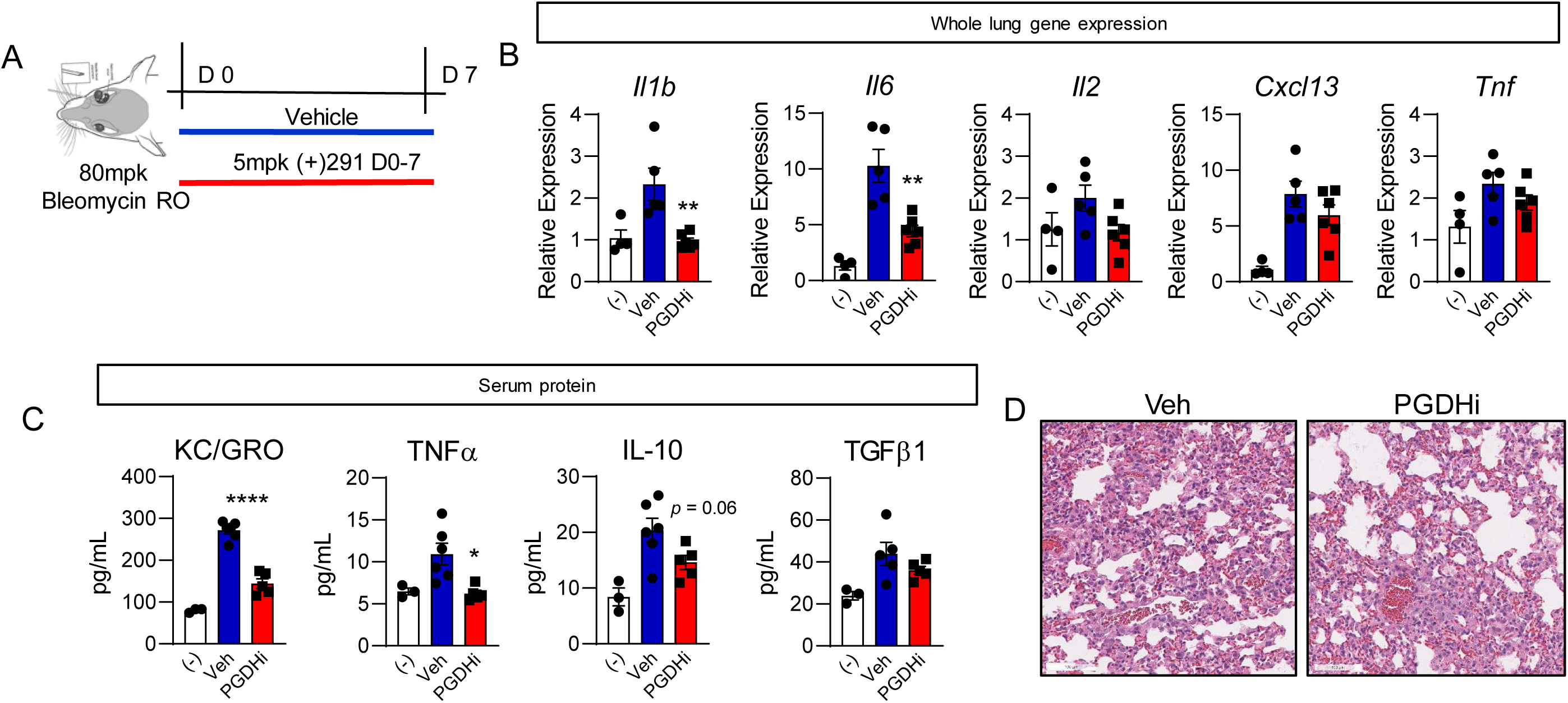
PGDHi mitigates bleomycin-induced inflammation. (**A**) Schematic depicting retro-orbital (RO) administration of 80mg/kg Bleomycin to 8wk old C57BL/6 female mice with subsequent PGDHi therapy (5mg/kg (+)SW033291, i.p. twice per day) and sacrifice at day 7. (**B**) Inflammatory factors *Il1b, Il6, Il2, Cxcl13, and Tnf* were quantified in lung homogenates isolated from naïve control mice (-), and vehicle (Veh) and PGDHi-treated mice 7 days post-bleomycin administration by RT-PCR and are expressed relative to *B2m* levels. (**C**) Inflammatory and fibrogenic factors KC/GRO, TNFα, IL-10, and TGFβ were measured in the serum of naïve (-), and Veh- and PGDHi-treated mice 7 days post-bleomycin administration by multiplex ELISA. Individual data points and mean±SEM are depicted; n=5-6 experimental mice/group. (**D**) Representative hematoxylin and eosin-stained lung sections from vehicle- and PGDHi-treated mice 7 days post-bleomycin administration. 20X, scale bars represent 100μm. ****P<0.0001, **P<0.005, *P<0.05 for PGDHi vs. Veh.

### PGDHi protects mice from severe weight loss, fibrosis, and lethality in bleomycin-induced IPF

The above results demonstrate that PGDHi mitigates acute inflammatory responses to bleomycin lung injury one week after exposure. To test whether PGDHi additionally protects against the severity of chronic IPF disease, we extended our analysis and continued to administer (+)SW033291 to bleomycin-exposed mice through day 35 (**Fig. 3A**). While vehicle-treated mice experienced severe weight loss in the first 8 days following bleomycin administration, PGDHi limited the severity and duration of weight loss. Specifically, vehicle-treated mice reached a nadir of 26.8% body weight loss on day 8 versus PGDHi-treated mice, which experienced only 16.0% maximum mean body weight loss on day 6 with weight stabilization thereafter (**Fig. 3B**). PGDHi-treated mice also achieved weight recovery closer to baseline between 21 and 28 days post-bleomycin administration, suggesting that PGDHi attenuated the effects of bleomycin on overall health. Notably, 33% of vehicle-treated mice succumbed to death, while the mortality rate of PGDHi-treated mice was less than 15% (**Fig. 3C**) Together these results suggest PGDHi therapy ameliorates systemic pathology during bleomycin-induced lung injury.

**Figure 3:**
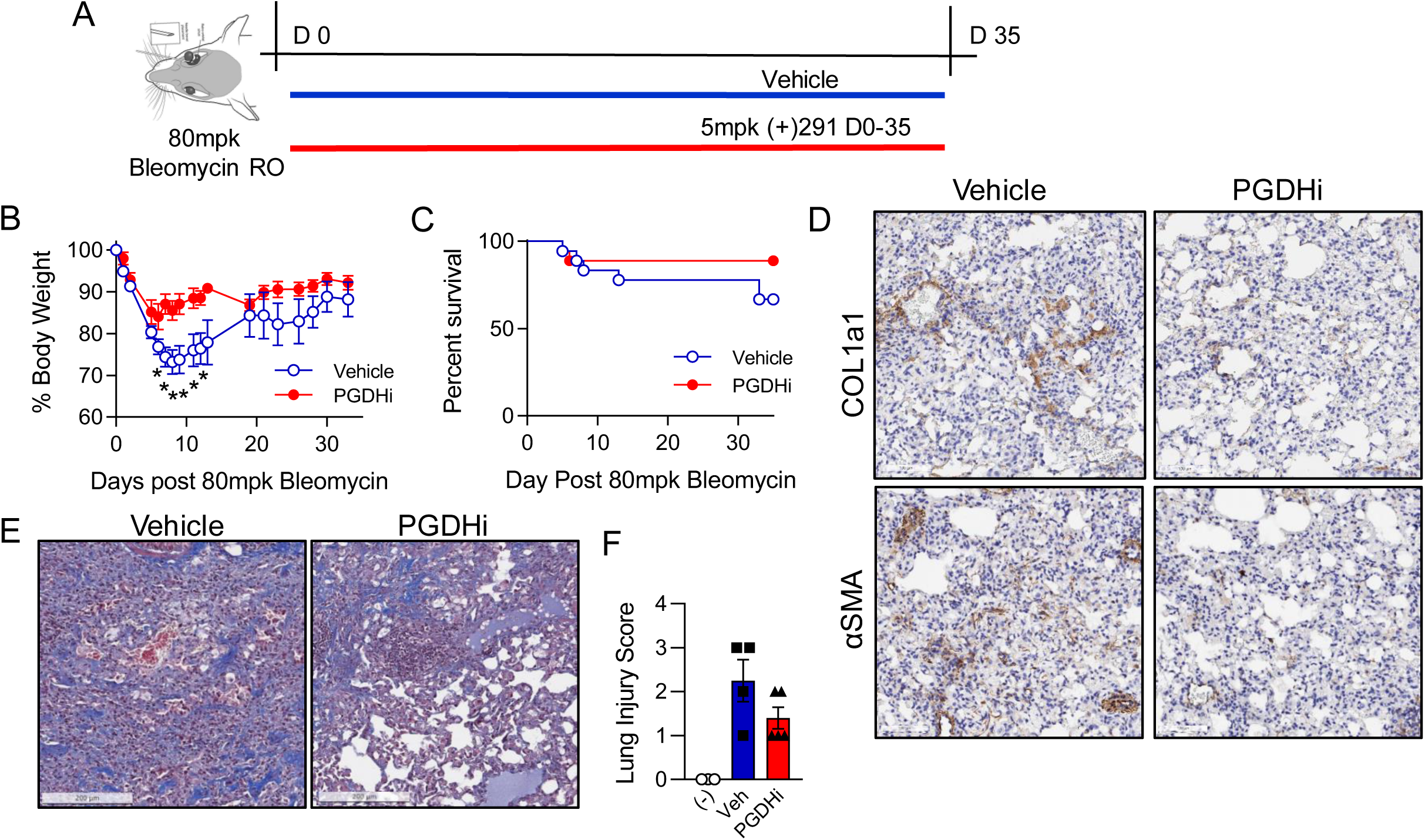
PGDHi mitigates Bleomycin-induced fibrosis. (**A**) Schematic depicting retroorbital (RO) administration of 80mg/kg Bleomycin to 8wk old C57BL/6 female mice with subsequent PGDHi therapy (5mg/kg (+)SW033291, twice per day) and sacrifice at day 35. (**B**) Body weight was measured for thirty-five days post-bleomycin administration as a percentage of day 0 weight for each mouse, treated as indicated. Mean±SEM is depicted. n=4-6 mice/group. (**C**) Kaplan-Meier survival curve following bleomycin administration. N=18 mice per group. (**D**) Representative images depicting COL1a1 and αSMA immunohistochemical staining (brown) in day 35 lung samples from vehicle- and PGDHi-treated mice. (**E**) Representative images of Masson’s trichrome-stained lung sections from vehicle- and PGDHi-treated mice 35 days post-bleomycin administration. (**F**) Pathological scoring of Masson’s trichrome-stained lung sections, based on a semi-quantitative morphological index of lung injury. Mean±SEM is depicted. n=4-5 mice per group. Scale bars represent 100μm. *P<0.05.

To determine the impact of PGDHi on bleomycin-induced lung fibrosis, ECM accumulation was evaluated in lung tissue 35 days post-bleomycin exposure. Importantly, pulmonary 15-PGDH expression was maintained following bleomycin administration (**Supplementary Figure**). Lung sections from vehicle-treated mice showed regional accumulation of Collagen 1a1 (COL1a1) and alpha-smooth muscle-actin (αSMA), whereas staining was reduced and non-focal in the lungs of PGDHi-treated mice (**Fig. 3D**), indicating that 15-PGDH inhibition protects against severe fibrosis in bleomycin-induced IPF. In addition to COL1a1, total collagen and lung morphology was evaluated on Masson’s trichrome-stained lung sections. Vehicle-treated mice displayed striking collagen deposition in the lung parenchyma, thickening of alveolar septa due to increased collagen, and intra-alveolar involvement. In contrast, PGDHi-treated mice showed fewer fibrotic lesions, which were interspersed with normal architecture, and less collagen accumulation (**Fig. 3E**). Pathological evaluation using a semiquantitiative morphological index of lung injury ^28^, where 0 represents normal lung tissue and 5 represents high levels of inflammation and fibrosis throughout the surveyed lung lobes, demonstrated substantial lung injury in vehicle-treated mice, which was attenuated by PGDHi (**Fig. 3F**).

### PGDHi attenuates pulmonary measurements of tissue stiffness in bleomycin-treated mice

To determine whether PGDHi-mediated reductions in inflammation and histopathological features of fibrosis corresponded to functional improvements, we measured static lung compliance (Cst) by generating pressure-volume loops in anesthetized, tracheostomized mice before necropsy. PGDHi treatment attenuated the impact of bleomycin on baseline static compliance, inspiratory capacity, and cellular content of the bronchoalveolar lavage fluid, as compared to vehicle-treated counterparts (**Fig. 4A**). Additionally, upon methacholine challenge, PGDHi-treatment resulted in significantly increased forced oscillation technique measurements of dynamic respiratory system compliance (Crs) and decreased respiratory system elastance (Ers; **Fig. 4B-C**). The bleomycin-induced loss in airway responsiveness to methacholine as Newtonian airway resistance (Rn), a measure of airflow in central airways, was also significantly reversed by PGDHi such that responses did not differ from naïve mice (**Fig. 4D**). Lastly, the rigidity of peripheral alveoli and lung parenchyma, measured as tissue elastance (H), was significantly reduced with PGDHi as compared to vehicle treatment (**Fig. 4E**).

**Figure 4:**
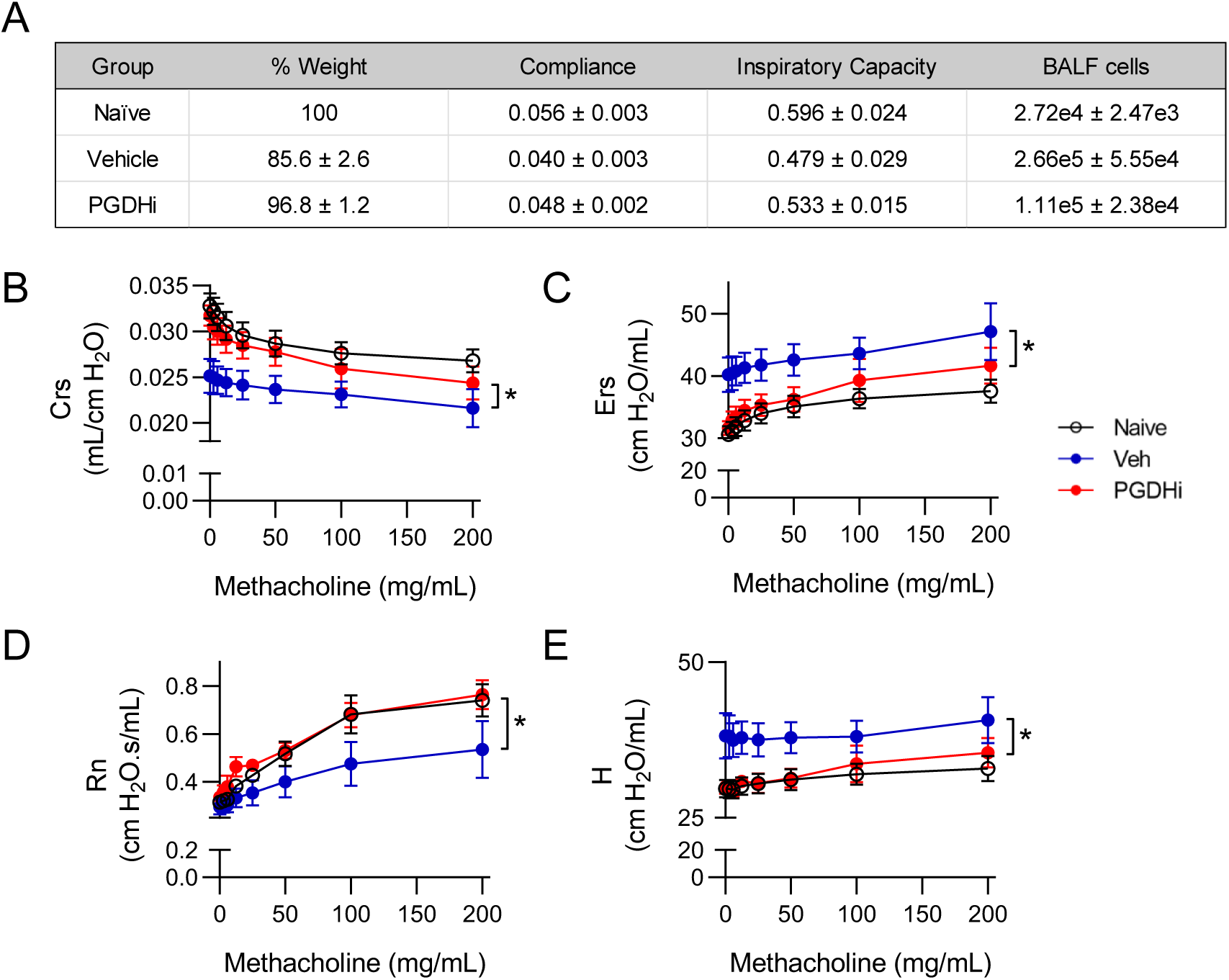
PGDHi mitigates the loss of pulmonary compliance and airway resistance observed in bleomycin-treated mice. PGDHi and vehicle treatment was administered through day 50 post-bleomycin administration. **(A)** Table depicting mean percentage of initial body weight, maximum baseline static lung compliance (mL per cm H2O), mean inspiratory capacity (mL), and mean number of cells collected in bronchoalveolar lavage fluid (BALF), ±SEM. n=3 mice/group. (**B-E**) Dynamic respiratory system compliance (Crs), respiratory system elastance (Ers), Newtonian airway resistance (Rn), and tissue elastance (H), measured using the forced oscillation technique at indicated doses of methacholine. Mean±SEM depicted. n=3 mice/group. *P<0.05.

### Endothelial cells, mast cells, and macrophages are PGDHi targets

Known cellular drivers of IPF pathogenesis include alveolar epithelial cells and fibroblasts ^1^. To identify the cell types that PGDHi may act directly on during IPF, we dissociated lung tissue from healthy mice and isolated cell populations based on surface marker expression. General hematopoietic and stromal separation on the basis of CD45 demonstrated enzyme activity in both CD45 positive and negative cells (data not shown). Stromal 15 -PGDH activity likely comprises endothelial cells as the CD31+ population demonstrated moderate activity enrichment. To further delineate the identity of the 15-PGDH+ hematopoietic cells, we fractionated on the basis of CD117 to enrich mast cells, which participate in pulmonary wound healing ^32^, and F4/80 to enrich alveolar macrophages, which are implicated in IPF pathogenesis^1^. Both CD117^+^ and F4/80^+^ preparations showed striking enzyme activity enrichment relative to negative counterpart cell fractions (**Fig. 5A**). Additionally, 15-PGDH staining colocalized with CD31, Toluidine blue, and F4/80 staining in serial sections (**Fig. 5B**). These data identify alveolar macrophages, mast cells, and to a lesser extent, endothelial cells, as the likely targets of PGDHi therapy in murine IPF.

**Figure 5:**
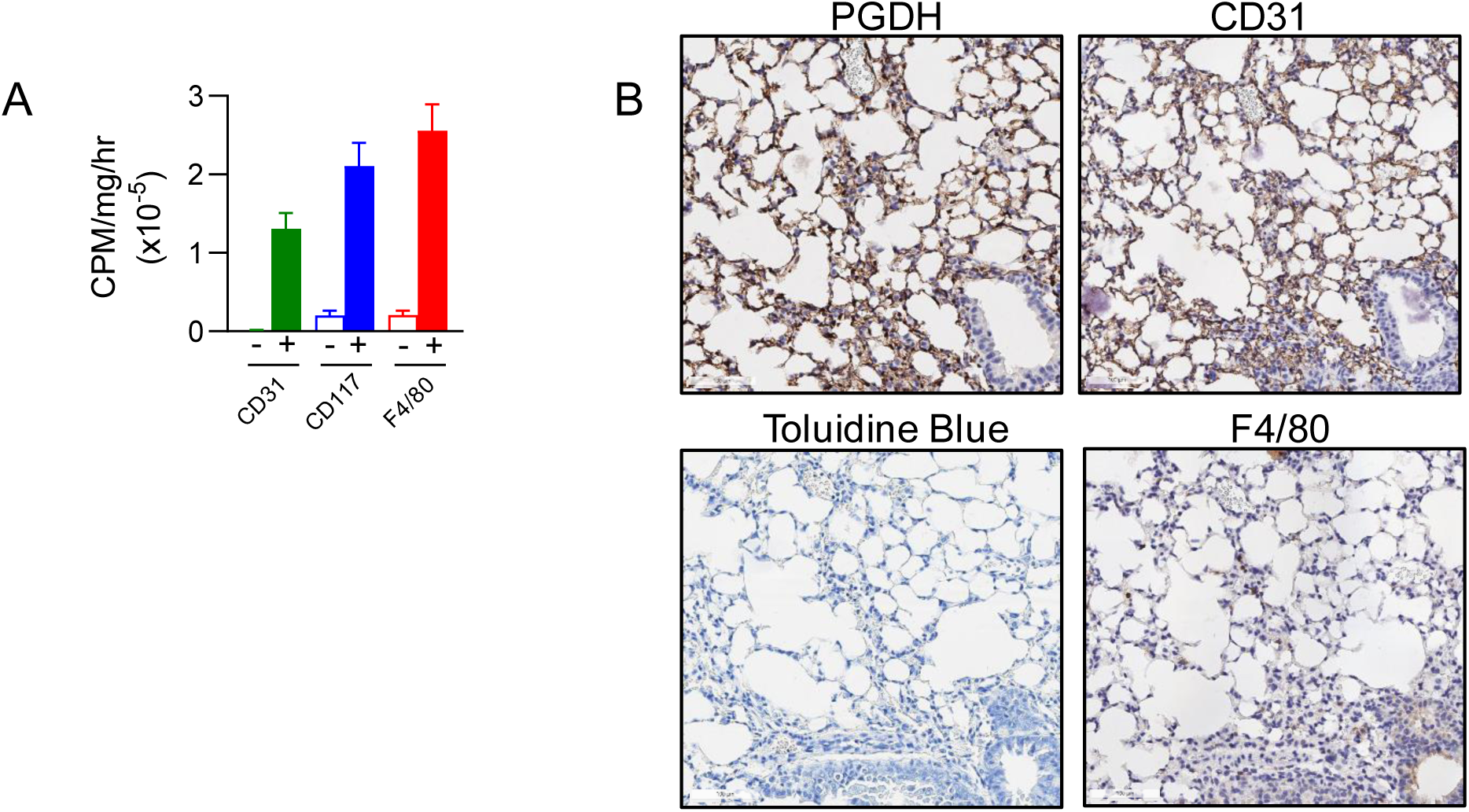
15-PGDH is enriched in murine endothelial, mast, and macrophage cells. (**A**) Quantification of 15-PGDH enzymatic activity in the indicated cellular populations (filled bars represent the positive fraction and open bars represent the negative fraction for each surface marker) isolated from the lungs of healthy mice, measured over one hour and normalized to input total protein (in mg). Mean±SEM is depicted. n=4-6 mice per group. (**B**) Representative images of 15-PGDH, CD31, Toluidine Blue, and F4/80 staining (all in brown) in serial sections of healthy murine lung (20X). Scale bar represents 100μm.

### PGDH+ cells are present in human lung

Taken together, our *in vivo* results strongly identify 15-PGDH as a therapeutic target in murine bleomycin-induced IPF. To determine if therapeutic targets exist in human lung tissue, we characterized 15-PGDH expression in human lung sections incidentally removed on autopsy. 15-PGDH was expressed in 6 of 6 lung specimens examined, and localized to cells lining the alveoli and blood vessels (**Fig. 6**), consistent with a previous report ^33^. The existence of PGDH+ cells in human lung tissue supports the notion that 15-PGDH inhibition may be a novel and effective therapy for IPF patients.

**Figure 6:**
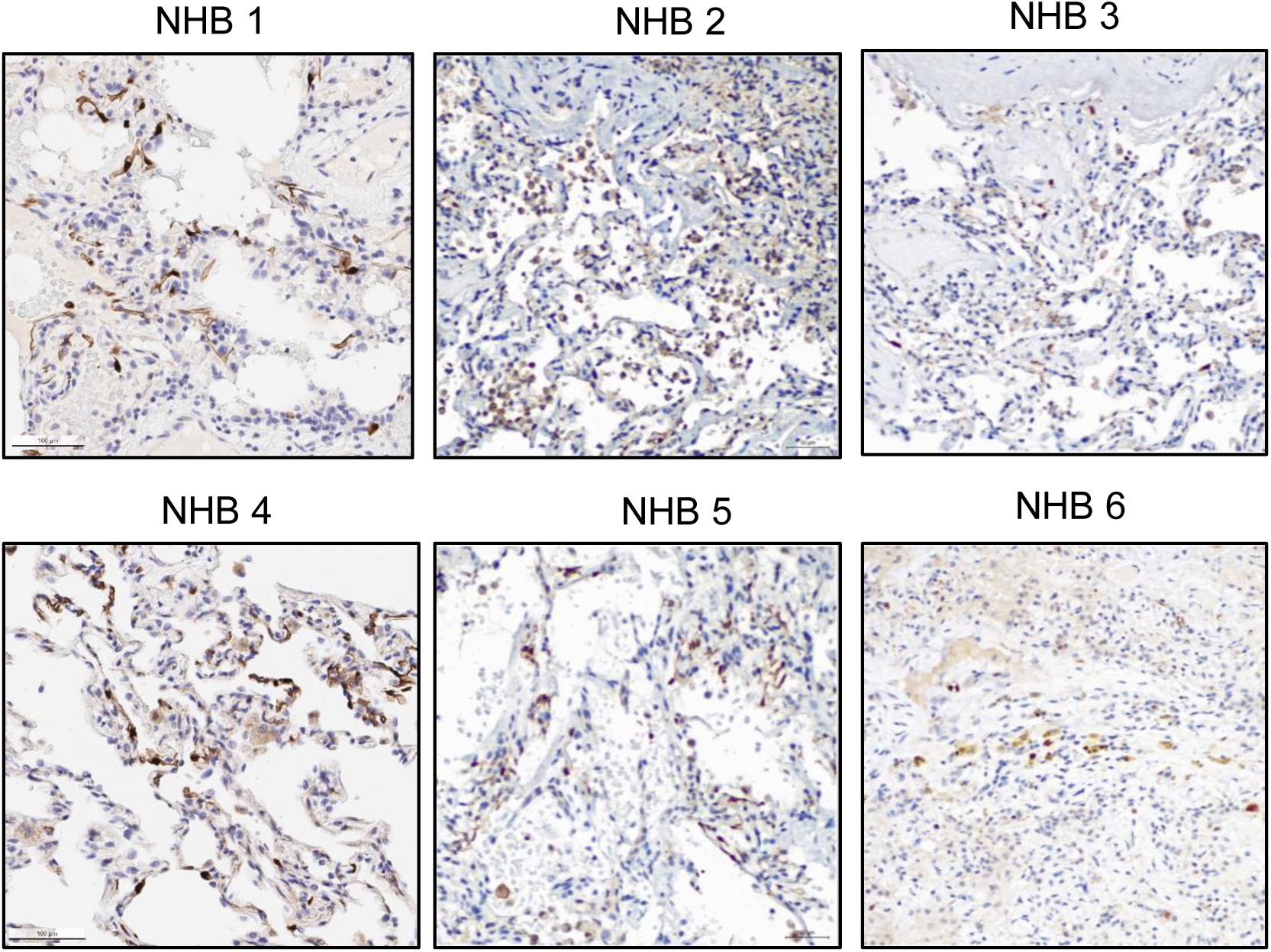
15-PGDH is expressed in human lung. Representative images of 15-PGDH staining (brown) in human lung (20X) from 6 unique biopsies. Scale bar represents 100μm.

## Discussion

In this study, we identified the presence of abundant 15-PGDH+ cells in murine pulmonary tissue. Compared to bone marrow and colon; tissues in which we have previously demonstrated enhanced regeneration and wound healing via pharmacologic 15-PGDH inhibition, lung tissue shows greater expression and enzyme activity. As PGE2 is well known to antagonize TGFβ-mediated fibrotic signaling, we tested the capacity for 15-PGDH inhibition to protect against IPF pathology in mice exposed to bleomycin. Our data show that PGDHi strikingly attenuates early inflammation at day 7, which correlated with reduced weight loss and mortality. We find that continuous PGDHi treatment reduces the occurrence of fibrotic lesions and interstitial remodeling in the lung at day 35. Moreover, PGDHi attenuated lung dysfunction, as measured by changes in compliance, resistance, and tissue rigidity. Our data identify pulmonary endothelial cells, macrophages, and mast cells as the direct cellular targets of PGDHi. Lastly, the presence of PGDH+ cells in human pulmonary tissue identifies 15-PGDH as a potential novel therapeutic target in human IPF.

These data extend previous work demonstrating the capacity for PGE2 or specific agonism of EP2/EP4 receptors to attenuate TGFβ-mediated pro-fibrotic signaling in pulmonary fibroblasts ^19,23-25^, and provide a well-tolerated and efficacious alternative to the delivery of long-acting prostaglandin analogs ^18^. Our dual therapeutic effect on both acute inflammation and chronic fibrosis, as well as, our finding that pulmonary endothelial cells, mast cells, and macrophages are highly 15-PGDH active, suggest multiple discrete mechanisms contribute to PGDHi-mediated protection. These results also indicate that systemic 15-PGDH inhibition may exert a more potent protective effect than targeted PGE2 delivery to endothelial cells ^21^ or alveolar macrophages ^22^ during IPF. Early in disease, PGDHi may improve alveolar epithelial wound repair, in a manner similar to that which has been demonstrated in the colonic epithelium^26^. The concept that PGDHi protects in part by improving alveolar stem and progenitor cell function is consistent with recent reports that senescent epithelial cells and fibroblasts drive IPF pathology ^34^. Later in disease, and of particular clinical relevance, PGDHi reduces ECM protein expression and fibrotic remodeling in the lung, consistent with the impact of PGE2 administration or specific EP2/EP4 agonism, on human pulmonary fibroblasts from IPF patients^19^. Notably, PGDHi was associated with fewer fibrotic foci, preservation of alveolar architecture, and striking improvements to the dynamic compliance, resistance, and parenchymal rigidity of lungs upon forced oscillation analysis.

During the preparation of this manuscript, corroborating data was published by Bärnthaler *et al*. ^35^. Both studies agree in the ability of PGDHi to increase local PGE2 levels, to prevent the accumulation of collagen, and to improve lung function in murine IPF. Our work additionally demonstrates robust PGDH enzymatic activity in pulmonary tissue as compared to additional organs in which PGDHi has therapeutic efficacy ^26^. Moreover, we demonstrate PGDHi-mediated amelioration of acute inflammatory pathology, including local diminution of Il1b and Il6, and systemic reductions in KC/GRO, and TNFα, which likely contribute to the subsequent reductions in fibrosis and improvements in pulmonary function.

PGDHi provides a novel and well-tolerated therapeutic approach to IPF, which is increasing in prevalence worldwide, particularly among aging populations. In the United States prevalence is estimated at 4.0 per 100,000 persons aged 18-34, and 227.2 per 100,000 persons among those >75 years ^36^. A history of cigarette smoking is the strongest risk factor, with additional risks including exposure to stone, wood, and organic dusts, and gastroesophageal reflux, which may contribute to lung injury via microaspiration ^37,38^. While recent work has yielded greater insight into the pathogenesis of IPF and related progressive fibrosis that occurs in interstitial pneumonias of unknown cause, treatment options remain only partially effective. FDA approved therapies include Nintedanib, an inhibitor of nonreceptor and receptor tyrosine kinases, and Perfenidone a small molecule of unknown mechanism. Neither are curative but demonstrate an ability to slow disease progression and can be combined with palliative care approaches to improve quality of life ^37,38^. Patients deemed critical may be eligible for lung transplantation, which provides a 40-50% 5-year survival rate ^39^. With a clear need for improved therapies for IPF and other interstitial lung diseases, our work demonstrates PGDHi holds significant promise either as a standalone therapy to prevent myofibroblast differentiation and reduce collagen deposition, or as a combination strategy with the currently approved therapies and palliative care.

Of interest, this work has additional therapeutic implications for disease states also characterized by fibrotic lesions. Primary myelofibrosis (PMF) is a rare bone marrow disorder characterized by abnormal blood cell production which results in extensive bone marrow scarring, activation of extramedullary hematopoiesis to the spleen, and a high incidence of leukemic transformation ^40^. Median overall survival is only six years using current standards of care ruxolitinib, hydroxyurea, or bone marrow transplantation. ^41-43^. PGDHi may hold significant promise for PMF therapy, as inhibition of the TGFβ signaling axis has demonstrated efficacy in preclinical PMF models ^44,45^. TGFβ transformation of fibroblasts plays a significant role in additional fibrotic diseases including hepatic, skin, and post-radiation induced fibrosis of multiple organs ^46^, as well. We propose that PGDHi-mediated disruption of myofibroblast differentiation will have therapeutic benefit in a number of these models and provide novel treatment strategies for these poor prognosis conditions.

## Methods

### Reagents

Bleomycin sulfate was purchased from Cayman Chemical and was dissolved in 0.9% sodium chloride to a concentration of 16mg/mL. 80mg bleomycin per kg mouse body weight was administered as a single dose at day 0 by retro-orbital injection under isoflurane anesthesia. (+)SW033291 was prepared in a vehicle of 10% ethanol, 5% Cremophor EL, 85% dextrose-5 water, at a concentration of 125ug/200ul for use at 5mg/kg for a 25g mouse. (+)SW033291 was administered by intraperitoneal injection, twice per day spaced by 8 hours, beginning immediately after bleomycin administration and continuing through the duration of the experiment.

### Animals

The animals described in this study were housed in the AAALAC accredited facilities of the CWRU School of Medicine. Standard Operating Procedures and reference materials are available from the IACUC Office for animal use. The animal health program was directed by the Case Animal Resource Center Director, W. John Durfee, DVM, Diplomate ACLAM, and provided by two full-time clinical veterinarians. Steady-state analysis and bleomycin administration was performed on female C57BL/6J mice obtained from Jackson Laboratories at 8 weeks of age. All animals were observed daily for signs of illness, and following bleomycin administration, mice were also weighed 2 to 3 times per week. Mice were housed in standard microisolator cages and maintained on a defined, irradiated diet and autoclaved water. Medical records and documentation of experimental use were maintained individually or by cage group. Veterinary technicians under the direction of the attending veterinarian provided routine veterinary medical care, if needed. Animal care and use was additionally monitored for training and compliance issues by the Director, Research Compliance IACUC. The Case Assurance number is A-3145-01, valid until April 30, 2023. All husbandry and experimental procedures were approved by the Case Western Reserve University Institutional Animal Care and Use Committee (IACUC) and we confirm that all procedures were performed in accordance with approved IACUC protocol 2013-0182.

### Histological and immunohistochemical analysis

Animals were harvested via CO2 inhalation followed by cervical dislocation and whole lungs excised and placed in 10% neutral buffered formalin for 24 hours. Samples were transferred to PBS and shipped to Histowiz where they were embedded in paraffin, and sectioned at 4μm. Immunohistochemistry was performed according to Histowiz protocols (https://home.histowiz.com/faq/). Histowiz defines their standard methods as the use of a Bond Rx autostainer (Leica Biosystems) with enzyme treatment using standard protocols, and detection via Bond Polymer Refine Detection (Leica Biosystems) according to manufacturer’s protocol. Whole slide scanning (40x) was performed on an Aperio AT2 (Leica Biosystems).

### Real Time PCR

On Day 0 and 7 post bleomycin induction, animals were harvested via CO2 inhalation followed by cervical dislocation and whole lungs excised and homogenized in RLT buffer. For comparison of bone marrow, colon, and lung, hindlimb marrow was flushed and the distal colon was dissected, prior to homogenization in RLT buffer. RNA isolation was performed using the RNAEasy Kit (Qiagen) and real-time PCR performed using commercial primers purchased from Applied Biosystems: IL2 (Mm00434256_m1), IL1B(Mm00434228_m1), IL6 (Mm00446190_m1), TNF (Mm00443258_m1), CXCL13 (Mm04214185_s1). Values were tabulated graphically with error bars corresponding to standard error of the means and compared using 2-tailed t-tests.

### Multiplex ELISA

Peripheral blood from mice 7 days post-bleomycin exposure was collected into Microtainer serum-separator tubes (Becton-Dickinson) by submandibular cheek puncture. Whole blood was allowed to clot at room temperature and then spun at 6000 x g for 3 minutes to separate serum. Serum was removed and stored at -80 prior to analyzing with the V-PLEX ProInflammatory Panel 1 Mouse Kit (Meso Scale Diagnostics).

### Lung Mechanics

Mice were weighed and anesthetized with an intraperitoneal injection of ketamine/xylazine and placed in a supine position. Once mice were nonresponsive to toe pinch, midline tracheostomy was performed to insert a blunt tip cannula. Mice were then paralyzed with pancuronium bromide and mechanically ventilated by the flexiVent system (SCIREQ, FX2 module) with continuous ECG monitoring. The maximum baseline static compliance was measured using computer software slope-fit of a quasi-static pressure-volume loop (Salazar-Knowles equation). Dynamic compliance (Crs), elastance (Ers), Newtonian airway resistance (Rn), and tissue elastance (H) were calculated by forced oscillation technique (Snapshot-150 and Quick Prime-3). Following two recruitment breaths (30 cm H_2_O for 3 seconds), increasing doses of methacholine (0-200 mg/mL; Sigma-Aldrich) were administered over 10 seconds via an in-line ultrasonic nebulizer (AeroNeb, SCIREQ). After each methacholine dose, 6 dynamic measurements were made and the mean values were analyzed. Subsequently, the bronchoalveolar lavage fluid was collected by instilling 0.5mL of PBS into the lung with a 10 second dwell time.

### Cell Separation

Briefly, lungs were minced and enzymatically-digested using previously described methods ^47^ to generate a single cell suspension. Cells were then isolated by surface marker expression using Miltenyi microbead kits and LS column separation. 15-PGDH enzymatic activity was measured in lysed cell fractions, or in homogenized whole organs, as previously reported ^26^. Activity was then normalized to input protein and were tabulated graphically with error bars corresponding to standard error of the means and compared using 2 - tailed t-tests.

### Statistical Analysis

Analysis was performed using GraphPad Prism software. Unpaired two-tailed Student’s t-test was used to compare between groups, unless otherwise noted. For body weight loss a two-way ANOVA with a post-hoc Sidak’s multiple comparisons test was used. For lung mechanics, area under the curve was calculated and unpaired two-tailed Student’s t-tests were used to compare vehicle versus PGDHi responses.

## Acknowledgments

This work was funded by NIH grants R35 CA197442 and K99 HL135740, and was supported by the Lung Infection and Inflammation Modeling Core the Tissue Resources Core Facility of the Case Comprehensive Cancer Center (P30CA043703).

## Author Contributions

J.N.P.S. conceived the hypothesis, designed the experiments, collected and interpreted the data, and wrote the manuscript. M.D.W. collected and interpreted the data. A.P.J. and K.F.C. collected the data. T.M.R. collected and interpreted the lung mechanics data and revised the manuscript. S.D.M. conceived the hypothesis and revised the manuscript. A.B.D. conceived the hypothesis, designed the experiments, wrote the manuscript, and provided final approval of the manuscript. All authors have read the complete manuscript and provided final approval.

## Additional Information

### Supplementary Information

One supplemental figure accompanies this paper

### Supplemental Figure Legend

**Pulmonary 15-PGDH expression is maintained following bleomycin exposure**. Representative images of 15-PGDH staining in lung sections from vehicle- and PGDHi-treated mice at days 7 and 35 post-bleomycin exposure.

