## Supplemental Figure for "Therapeutic Targeting of 15-PGDH in Murine Idiopathic Pulmonary Fibrosis"

### Supplementary Figure: Pulmonary 15-PGDH expression is maintained following bleomycin exposure

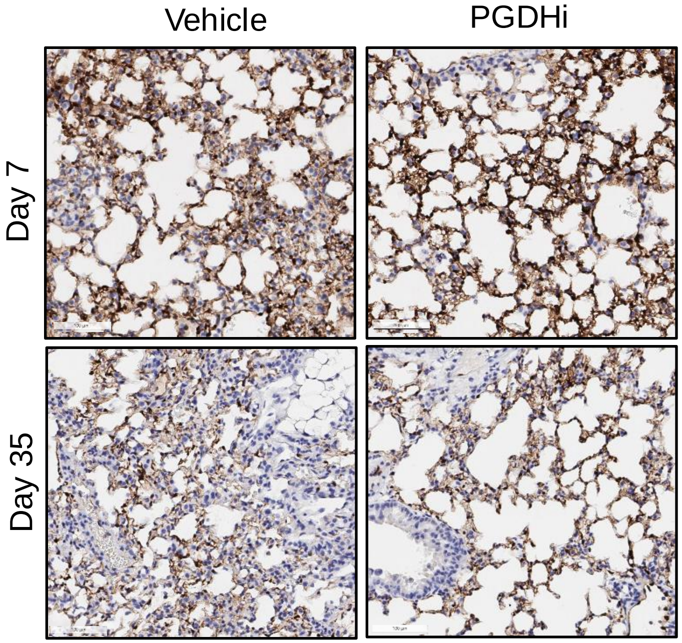


**Supplementary Figure 1: Pulmonary 15-PGDH is maintained following bleomycin.** Representative images of 15-PGDH staining in lung sections from vehicle- and PGDHi-treated mice at days 7 and 35 post-bleomycin exposure.
